# Function of Pumilio Genes in Human Embryonic Stem Cells and Their Effect in Stemness and Cardiomyogenesis

**DOI:** 10.1101/751537

**Authors:** Isabelle Leticia Zaboroski Silva, Anny Waloski Robert, Guillermo Cabrera Cabo, Lucia Spangenberg, Marco Augusto Stimamiglio, Bruno Dallagiovanna, Daniela Fiori Gradia, Patrícia Shigunov

## Abstract

Posttranscriptional regulation plays a fundamental role in the biology of embryonic stem cells (ESCs). Many studies have demonstrated that multiple mRNAs are coregulated by one or more RNA binding proteins (RBPs) that orchestrate the expression of these molecules. A family of RBPs, known as PUF (Pumilio-FBF), is highly conserved among species and has been associated with the undifferentiated and differentiated states of different cell lines. In humans, two homologs of the PUF family have been found: Pumilio 1 (PUM1) and Pumilio 2 (PUM2). To understand the role of these proteins in human ESCs (hESCs), we first demonstrated the influence of the silencing of PUM1 and PUM2 on pluripotency genes. *OCT4* and *NANOG* mRNA levels decreased significantly with the knockdown of Pumilio, suggesting that PUMILIO proteins play a role in the maintenance of pluripotency in hESCs. Furthermore, we observed that the hESCs silenced for PUM1 and 2 exhibited an improvement in efficiency of *in vitro* cardiomyogenic differentiation. Using *in silico* analysis, we identified mRNA targets of PUM1 and PUM2 expressed during cardiomyogenesis. With the reduction of PUM1 and 2, these target mRNAs would be active and could be involved in the progression of cardiomyogenesis.

## INTRODUCTION

Human embryonic stem cells (hESCs) are pluripotent cells derived from the inner cell mass of the blastocyst that have potential for differentiation into three germ layers (1–3). In an undifferentiated state, hESCs are characterized by the expression of stemness factors such as OCT4 (POU5F1), SOX2 and NANOG (4). These three transcription factors, which are positively regulated, are responsible for pluripotency maintenance and contribute to the repression of lineage-specific genes (reviewed by 5). When hESCs are stimulated to initiate the differentiation process, expression of genes associated with pluripotency is negatively regulated and genes associated with the germ layer begin to be positively regulated (6).

A complex network of gene expression underlies the molecular signaling that will give rise to the adult heart. Cardiomyogenic differentiation is a highly regulated process that depends on the fine regulation of gene expression (7). *In vitro* cardiomyogenic differentiation of hESCs mimics embryonic development and can be used as a model for cardiac development studies per se and as a model for research ranging from tissue electrophysiology to drug screening (reviewed by 8).

RNA-binding proteins (RBPs) are proteins that have RNA-binding domains and form ribonucleoprotein complexes in association with RNAs (RNPs). These proteins play a critical role in the posttranscriptional regulation of gene expression. The dynamics and function of these complexes depend on their composition, targets and cofactors (9). The PUF (Pumilio-FBF) family of RBPs is highly conserved among species and is found in plants, insects, nematodes and mammals (10–15). PUF proteins have RNA binding domains known as the Pumilio Homology Domain (PUM-HD). The Pumilio RNA interaction domain is highly conserved (16), comprising eight repeats, each having the ability to bind a single nucleotide of specific recognition motif in the 3′ untranslated region (UTR) of a target mRNA (17). In humans, two homologs of the PUF family are found: PUMILIO 1 (PUM1) and PUMILIO 2 (PUM2) which have 91% identity at the RNA binding domain (15).

The expression of PUM1 and PUM2 has been observed in hESCs and several human fetal and adult tissues, indicating a possible participation in the maintenance of germ cells (11,12). Furthermore, in mammals, the disruption of PUM proteins promotes defective germline phenotypes (18,19). In rodent, PUM1 facilitates the exit from the primitive state to the differentiated form by accelerating the degradation of some important elements in the maintenance of pluripotency, such as Tfcp2l1, Sox2, Tbx3, and Esrrb (20). Besides that, many of the mRNAs associated with PUM1 belong to a relatively small number of functional groups, suggesting an RNA regulon model (21) in which PUM1 inhibits translation and promotes the degradation of its target mRNAs (22). Pumilio proteins form multiprotein complexes with other regulatory proteins, such as DAZ-Like (DAZL) (11), BOULE (BOL) (23), Staufen (STAU) (24) and Nanos (NOS) (25,26). These complexes also are involved in the maintenance of ESC and in the regulation of the onset of meiosis in various organisms, including humans (11,27,28). PUM2 and NOS interact in a conserved mechanism for the development and maintenance of germ cells (29). Thus, the molecular scenario in which Pumilio proteins and their targets work, may lead to cellular differentiation or the maintenance of a stemness phenotype.

Here, we investigated the role of PUM1 and PUM2 proteins in hESCs during cardiomyogenic differentiation. We report that the silencing of PUM1 and PUM2 reduces the expression of pluripotency genes in hESCs. We found that the silencing of PUM1 and PUM2 positively influences cardiomyogenesis in hESCs. Using *in silico* analysis, we followed the targets of PUM1 and PUM2 during cardiomyogenesis, clustering the targets with the same behavior during differentiation and traced the gene networks.

## METHODS

### Cell culture and cardiac differentiation

The *NKX2-5*^*eGFP/w*^ HES3 hESC line (30) was obtained from Monash University (Victoria, Australia). The cell cultures were maintained on irradiated mouse embryonic fibroblasts (iMEFs) in hESC medium consisting of Dulbecco’s modified Eagle’s medium (DMEM)/F12 supplemented with 20% KnockOut serum replacement, 1% nonessential amino acids, 1% L-glutamine, 1% penicillin/streptomycin, 0.1 mM β-mercaptoethanol and 10 ng/ml human βFGF. The cells were passaged every 3–4 days by enzymatic dissociation using 0.25% trypsin/EDTA. Cardiomyogenic differentiation assays were conducted using an embryoid body (EB) protocol adapted from previously described (31,32) or a monolayer protocol previously reported (33).

Regarding EB cardiac differentiation protocol, briefly, 7×10^5^ cells/well were plated onto Growth Factor Reduced Matrigel® Matrix (Corning) 6-well coated dishes. Then, hESCs were dissociated and cultured on low-attachment plates with supplemented StemPro-34 medium (composed of StemPro™-34 SFM (Gibco™) supplemented with transferrin, ascorbic acid, penicillin/streptomycin and monothioglycerol) containing BMP4 (0.5 ng/mL) for 24 hours to form EBs (Day 0, D0). On day 1 (D1), the EBs were incubated with supplemented StemPro-34 added with βFGF (5 ng/ml), activin A (6 ng/ml) and BMP4 (10 ng/ml) to induce mesoderm differentiation. On day 4 (D4), the EBs were incubated with medium supplemented with XAV939 (10 µM/ml) and VEGF (10 ng/ml) to induce the cells to become cardiac progenitors. On days 8 and 11, the medium was changed to supplemented StemPro-34 containing only VEGF (10 ng/ml). On the 9^th^ day (D9), we observed the expression of eGFP. On day 15 (D15), we evaluated the efficiency of the protocol with cardiac troponin T (cTnT) staining. During the protocol cultivation period, from D0 to D15, the EBs were maintained in a humidified incubator under hypoxic conditions (5% O_2_, 5% CO_2_, 37°C).

For the monolayer cardiac differentiation protocol, hESC were dissociated from iMEF cultures and 1.5×10^5^ cells/well were plated onto Matrigel^®^ hESC-qualified Matrix (Corning) 24-well coated dishes. hESC were maintained in hESC medium until reached 90-100% of confluence to started the protocol. At day 0, RPMI medium supplemented with B27 without insulin (RPMI+B27-insulin) and 12 µM of CHIR99021 (Stemgent) were added to the culture. After 24h, medium was changed for RPMI+B27-insulin. At day 3, it was added RPMI+B27-insulin and 10 µM of XAV939 (Sigma) to the monolayer cultures, which were maintained until day 5, when medium was exchange for the RPMI+B27-insulin. From the 7^th^, the cultures were maintained with RPMI supplemented with B27 complete, with medium exchange every 3 days until the 15^th^. At final day (day 15), cells were fixed with paraformaldehyde 4% and stained for cTnT using a previously described immunofluorescence protocol (7). Analysis was performed on Operetta CLS High-Content Analysis System and Harmony software (PerkinElmer) (Supplementary Figure S1A).

### Lentiviral vector production and transduction

HEK293FT cells were cultured in Petri dishes containing DMEM supplemented with 10% fetal bovine serum, 1% L-glutamine and 1% penicillin and streptomycin for 24 hours. MISSION Lentiviral Mix and p-KLO1 containing short hairpin RNA (shRNA) targeting PUM1 (shPUM1), PUM2 (shPUM2) or a Scrambled sequence (shSc) (34) were added to the cell culture in OptiMEM containing Lipofectamine 2000 for 4 hours. Then, the medium was replaced with supplemented DMEM, as described above. After 48 and 72 hours, the medium was collected and centrifuged twice at 141000 *x g*. The cell pellet was resuspended in 1X PBS and stored at −80 ºC.

For transduction, hESCs were cultured on 6-well plates and different dilutions of the lentiviruses were tested (Supplementary Figure 2), and the dilution was defined as 10^−3^ for all experiments. After the transduction, the medium was replaced and hESCs were cultured for 24 hours to induce cardiac differentiation.

### RNA extraction and quantitative RT-qPCR

RNA was extracted using the RNeasy Kit (Qiagen), and the cDNA reaction was performed using the Improm II Kit (Promega) according to the manufacturer’s instructions. Samples were obtained from three replicates for undifferentiated cells and from three independent cardiac differentiations. Amplifications were carried out in a final volume of 10 μl containing SYBR Select master mix (Applied Biosystems), 100 ng cDNA template, and 5–10 pmol primers. The RT-qPCR conditions followed the manufacture’s recommendations (Applied Biosystems), using the LightCycler system (Roche). The melting curves were acquired after RT-qPCR to confirm the specificity of the amplified products (Supplementary Table S1). We generated standard curves for each gene, including the GAPDH (housekeeping) gene. Amplifications were performed in triplicate. Cq results for each gene were normalized based on GAPDH expression and the analysis of relative expression were performed.

### Flow cytometry

During cardiac differentiation, the cells were immunophenotyped by flow cytometry to confirm differentiation stages. The EBs were dissociated with 0.25% trypsin/EDTA (5 min) and resuspended in PBS/0.5% BSA. On day 3, the cells were incubated with anti-CD56 (BD) antibody (1:12.5) for 20 min at 4 ºC. On day 9, the cells were only dissociated for eGFP detection. On day 15, for cTnT staining, the EBs were incubated with trypsin/0.25% EDTA for 20 min, followed by inactivation with DMEM supplemented with 50% SFB and DNase I (20-30 U/ml). After dissociation, the cells were fixed with 4% formaldehyde (20 min), followed by permeabilization with 0.5% Triton X-100 (25 min). Then, the cells were incubated with an anti-cTnT primary antibody (Thermo Fisher Scientific) (1:100) for 30 min at room temperature, followed by incubation with Alexa Fluor 633 secondary antibody (1:1000) (30 min). The cells were analyzed on a FACSCanto II (BD) flow cytometer. Data analysis was performed using FlowJo software (v.10).

### Immunofluorescence

The immunofluorescence protocol followed as previously described (7). Briefly, monolayer cultures we fixed with paraformaldehyde 4%, rinsed with PBS, followed by incubation with blocking buffer (PSA/BSA 5%) for 60 min. Then, cells were incubated with primary antibodies for Pumilio 1 (1:300, Bethyl laboratories Inc.), Pumilio 2 (1:70, Bethyl laboratories Inc.) or OCT4 (1:100, Abcam), diluted in blocking buffer, for 60 min, at 30°C. After three PBS washes, cells were incubated with Alexa Fluor® 488 anti-goat or anti-rabbit secondary antibody, for 60 min, at 30°C. DAPI were staining for 10 min, followed by three PBS washes. Analysis was performed on Operetta CLS High-Content Analysis System and Harmony software (PerkinElmer) (Supplementary Figure S1B).

### Analysis of mRNA targets of PUM1 and PUM2

To analyze the mRNA targets of PUM1 and PUM2 proteins through cardiac differentiation, we used the RNA-seq data from mRNAs associated with polysomes obtained during in vitro cardiomyogenesis previously published (32). Initially we defined a set of 1,809 target genes for human Pumilios from published (22). Differential expression analysis was done using the Bioconductor R package edgeR (35,36). Comparisons were performed for polysome-bound RNA fractions - each sample against the preceding time-point: Day 0 vs Day 1 and Day 0 vs Day4. For these analyses, we retained only those genes with at least one count per million in at least three samples. Based on initial results of DEGs we used a stringent analysis using a p-value threshold of 0.05 and a log2Fold change (logFC). Genes with a logFC > 1 were considered upregulated, and genes with a logFC < −1 were considered downregulated. An enrichment analysis of this set of genes was performed using g:Profiler (37) (http://biit.cs.ut.ee/gprofiler/) and REVIGO (38) (http://revigo.irb.hr/) consortium database. Complement analysis was performed with STRING CONSORTIUM 2019 (39) (https://string-db.org).

### Statistical analysis

Statistical analysis was performed using GraphPad Prism 7 software. The data sets are expressed as the means ± standard deviation. According to data sets were used unpaired Student’s t-test or one-way ANOVA followed by Tukey post hoc test. Differences with p<0.05 were considered statistically significant.

## RESULTS

### Differences in *PUM1* and *PUM2* expression comparing total and polysomal RNA

To evaluate the expression levels of *PUM1* and *PUM2* throughout cardiomyogenesis, hESCs were submitted to an *in vitro* cardiac differentiation protocol and its progression was verified by flow cytometry (Figure 1A). On D3, about 25% of the cell population was CD56^+^ (mesoderm marker) indicating an advanced process of mesodermal differentiation (Figure 1B). On D9, the expression of eGFP, under the control of the NKX2.5 promoter (cardiac progenitor marker), was evaluated and approximately 20% of the cell population expressed the marker, confirming the cardiac commitment of this population. On the 15^th^ day, the expression of the cardiomyocyte marker cTnT was assessed, showing positive labeling for approximately 20% of the cell population (Figure 1B).

**Figure 1.**
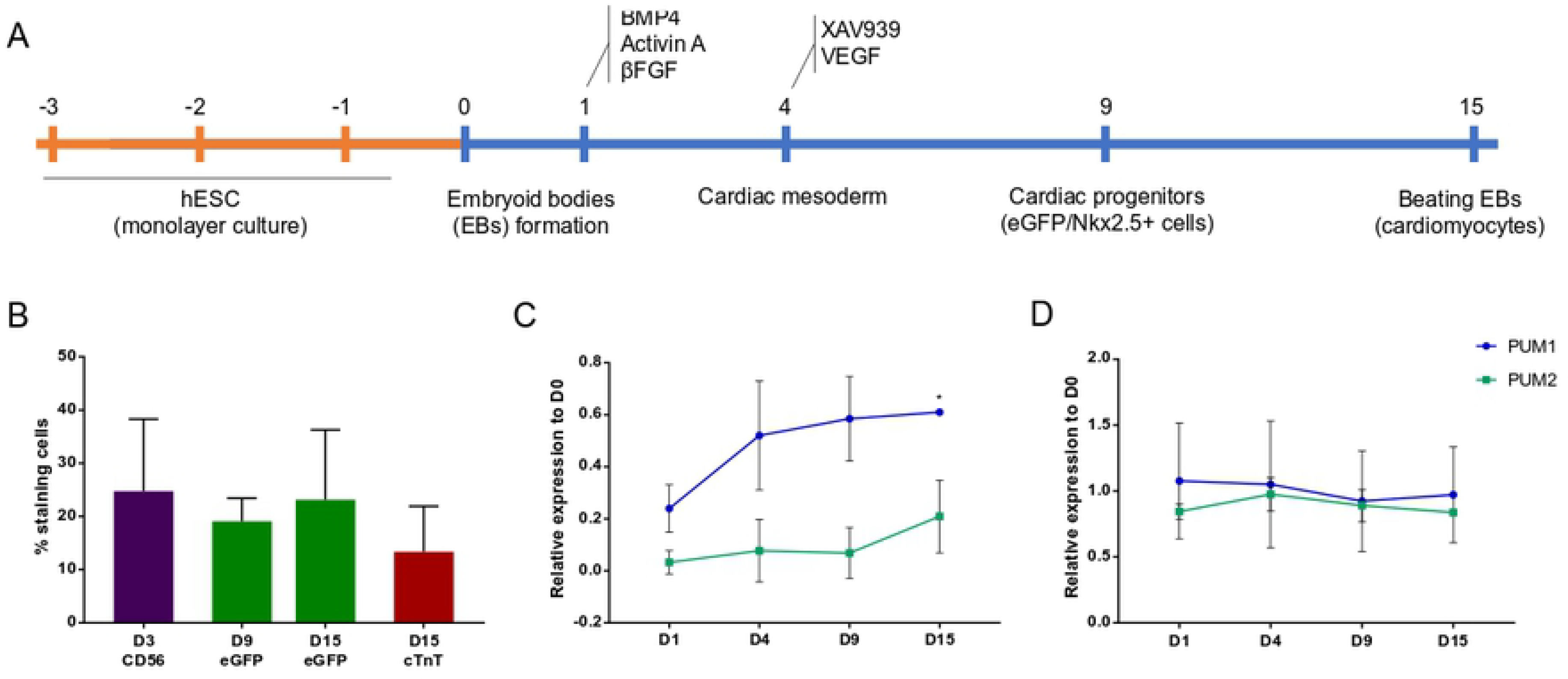
*PUM1* and *PUM2* expression profile during cardiomyogenesis of hESC. (A) Scheme of the EB cardiomyogenic differentiation protocol of hESC. (B) Graph indicating the percentage of positive cells for the markers CD56, eGFP/Nkx2.5 and cTnT in different time-points of in vitro cardiac differentiation. (C) Relative expression of *PUM1* and *PUM2* during days 1, 4, 9 and 15 of cardiomyogenesis, in relation to day 0 (total RNA). (D) Relative expression of RPKM values of *PUM1* and *PUM2* mRNAs associated with polysomes during in vitro cardiomyogenesis, in relation to day 0 (Data extracted from previously published results by 31). *p<0.05 in relation to D1.

Total RNA was extracted from 5 time-points during cardiac differentiation (D0, D1, D4, D9, D15) and RT-qPCR was performed. The expression analysis of *PUM1* and *PUM2* mRNAs showed a slight increase throughout the differentiation process (Figure 1C). In previous work, we analyzed the expression of polysome-associated mRNAs along cardiomiogenesis (32). Interestingly, when we analyzed the association of PUM mRNAs with polysomes along cardiomyogenesis, based on the fold change of Reads Per Kilobase Million (RPKM) values comparing each time-point in relation to D0, we noticed no significant changes (Figure 1D).

### Individual and combined knockdown of *PUM1* and *PUM2* affects the expression of *OCT4* and *NANOG* in hESCs

To understand the role of PUM1 and PUM2 in hESC maintenance or during cardiomyogenic differentiation, we silenced their expression using short hairpin RNAs. We produced lentiviral particles containing shRNA that recognize *PUM1*, *PUM2* and a scrambled control from previously published plasmids (34) (Supplementary Figure S2). After 24 h of hESCs transduction, the cells showed no evident morphological changes (Figure 2A). Knockdown of *PUM1* and *PUM2* was confirmed by RT-qPCR, which showed a significant reduction of *PUM1* and *PUM2* mRNA levels under silencing conditions compared to the control in single- and double-silenced cells (Figures 2B-C). sh*PUM1* also interfered in reducing the expression levels of *PUM2* mRNA and vice versa, suggesting that the shRNAs presented a low specificity or the PUM proteins could be regulating each other. Double-silencing showed better efficiency in silencing both Pumilio genes: 92% for *PUM1* and 90% for *PUM2*.

**Figure 2.**
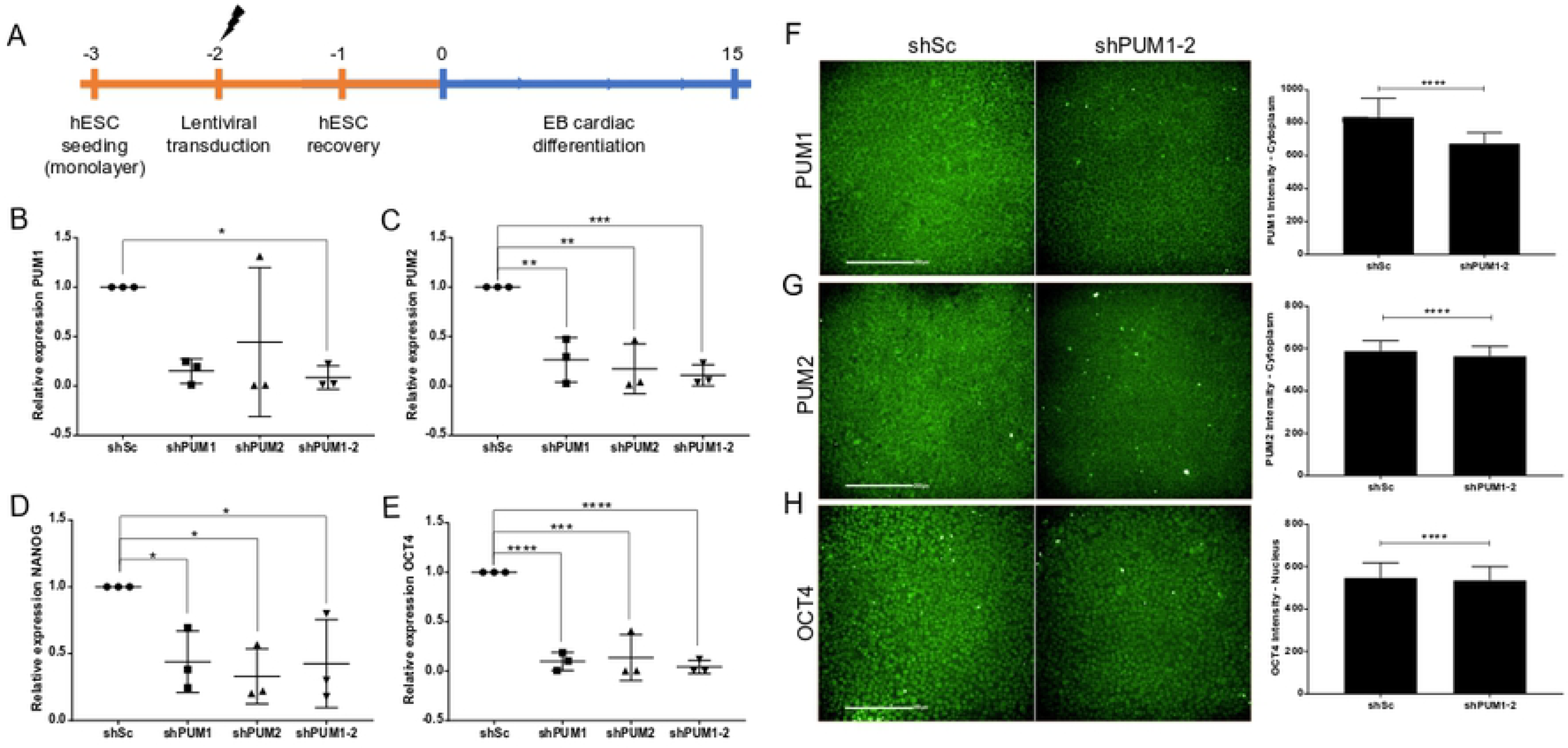
Knockdown of *PUM1* and *PUM2* affects the expression of *OCT4* and *NANOG*. (A) Transduction scheme of hESC before EBs cardiac differentiation. (B-E) Analysis of the relative expression of *PUM1* (B), *PUM2* (C)*, NANOG* (D) and *OCT4* (E) in hESCs after 24 h of transduction with lentiviral vectors containing shSc, shPUM1, shPUM2 and shPUM1-2. Representative images of immunostaining and its respective intensity quantification of *PUM1* (F), *PUM2* (G) and *OCT4* (H) after 24 h of transduction with lentiviral vectors containing shSc and shPUM1-2. Scale bars: 200 μm. *p<0.05, **p<0.01, ***p<0.001, ****p<0.0001

Then we observed whether the silencing of *PUM1* and *PUM2* altered the mRNA levels of some pluripotency transcription factors, as *OCT4* and *NANOG*. Interestingly, the mRNA levels were significantly reduced for both genes in cells with *PUM1* or *PUM2* silencing and double silencing relative to the control (Figures 2D-E). Considering the efficacy of shPUM1-2 in reducing Pumilios and OCT4 mRNA expression, we quantified the protein content. Quantification of cytoplasmatic *PUM1* and *PUM2* and nuclear *OCT4* staining intensity comparing hESC cultures transduced with shPUM1-2 with those transduced with shSc (Figure 2F-H) indicated a decrease in protein quantity. These results suggest that PUM1 and PUM2 could regulate positively of *OCT4* and *NANOG* mRNAs.

### Knockdown of PUM1 and PUM2 in hESCs affects cardiomyogenesis

Considering the effects of Pumilio silencing in pluripotency factors, we also evaluated their influence in cardiomyogenesis. After 48h of hESC transduction, cardiac differentiation was induced. The transduction did affect neither the morphology of cells nor the formation of EBs (D1 and D4) (Figure 3A). From the 9^th^ day, silenced EBs, visually, presented some morphological differences with respect to their size compared to the control cells (Figure 3A), though this difference was not statistically significant (Figure 3B). In all conditions, the EBs contract spontaneously (Supplementary Video 1).

**Figure 3.**
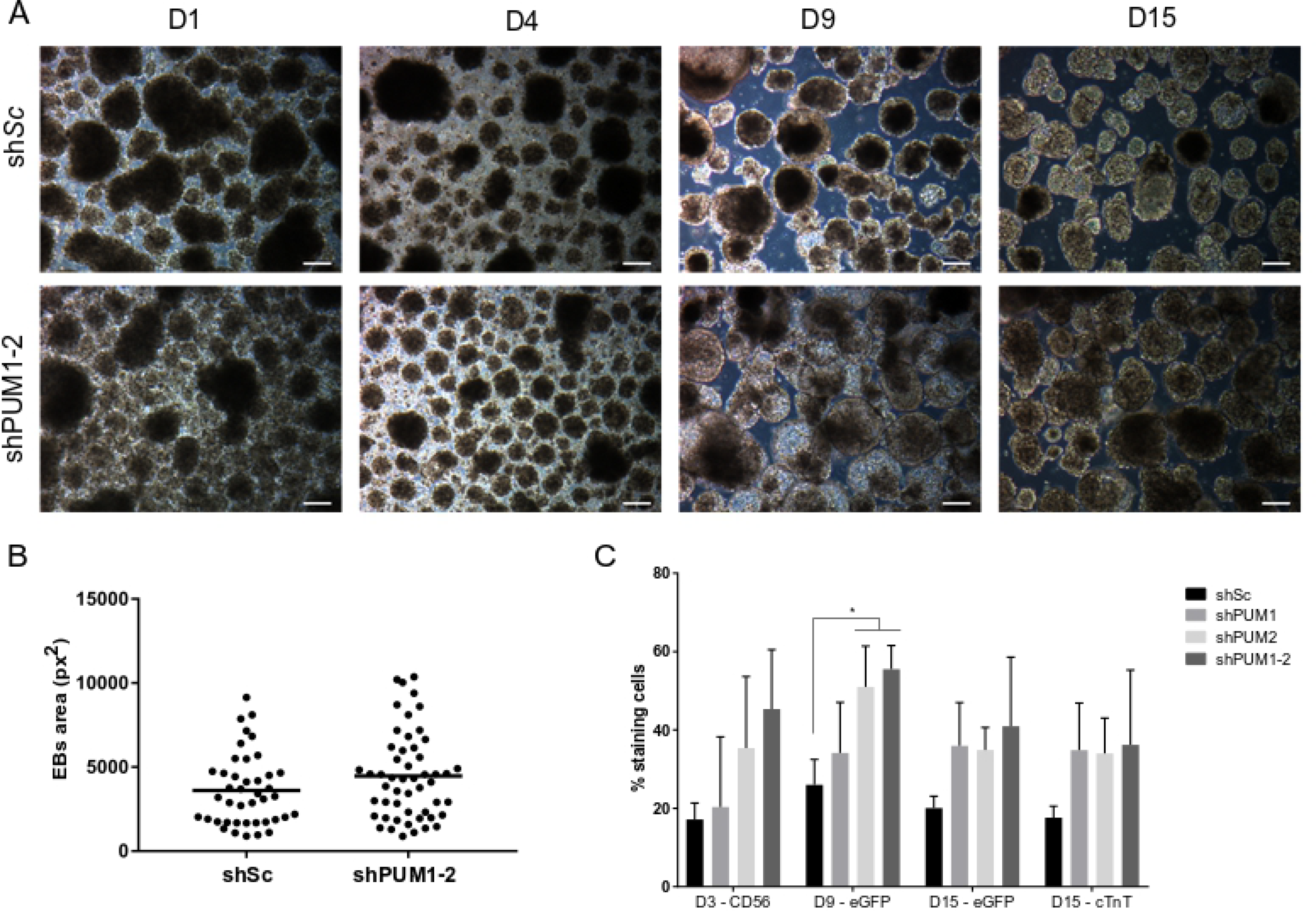
Effect of knockdown of *PUM1* and *PUM2* during EBs cardiac differentiation. (A) Morphology of EBs at days 1 (D1), 4 (D4), 9 (D9) and 15 (D15) of cardiomyogenesis differentiation of hESC previously transduced with shSc and shPUM1-2. Scale bars: 100 μm. (B) Area of EBs after 9 days of in vitro cardiac differentiation. (C) Percentage of positive cells for CD56, eGFP/Nkx2.5 and cTnT during cardiomyogenesis of the cells silenced for scramble, *PUM1*, *PUM2* and *PUM1-2*. *p<0.05

Analysis comparing shSc with shPUM knockdown conditions (shPUM1, shPUM2 and shPUM1-2) showed that cells transduced with shPUM2 and shPUM1-2 increase the number of eGFP+ cells at D9 compared to cells transduced with shSc, but not at D15 (Figure 3C). Regarding the efficiency of differentiation, on D15, the percentage of cTnT+ cells not changed between the different treatments (Figure 3C). These results demonstrated that when PUM was silenced, hESCs followed EBs cardiac differentiation efficiently, with no statistically significant changes.

We performed a monolayer cardiomyogenic differentiation protocol, as previously described (33) (Figure 4A). In this protocol we transfected hESCs with shSc or shPUM1-2 and analyzed the efficiency of cardiac differentiation through eGFP/Nkx2.5 expression and cTnT immunostaining at day 15. During the differentiation process, it was observed that *PUM1-2* silenced hESC had more beating areas in the monolayer culture than the shSc transduced cells, which were maintained until the end of the protocol (Supplementary Video 2). The beating frequency of shPUM1-2 transduced cells is on average 7 (±2.9) contractions in 10 seconds while shSc transduced cells showed an average of 3 (±1.3) contractions in 10 seconds (n=3, p<0,01) (Supplementary Video 2). Thus, the silencing of PUM1-2 increases the frequency of contractions relative to the shSc control. Also, the eGFP^+^ and cTnT^+^ stained areas, at the end of protocol, were significantly higher in shPUM1-2 treated cells, with no changes in cell number (DAPI^+^ area) (Figure 4B-D). Our data suggest that *PUM1* and *PUM2* could play a negative role in the control of cardiac lineage commitment.

**Figure 4.**
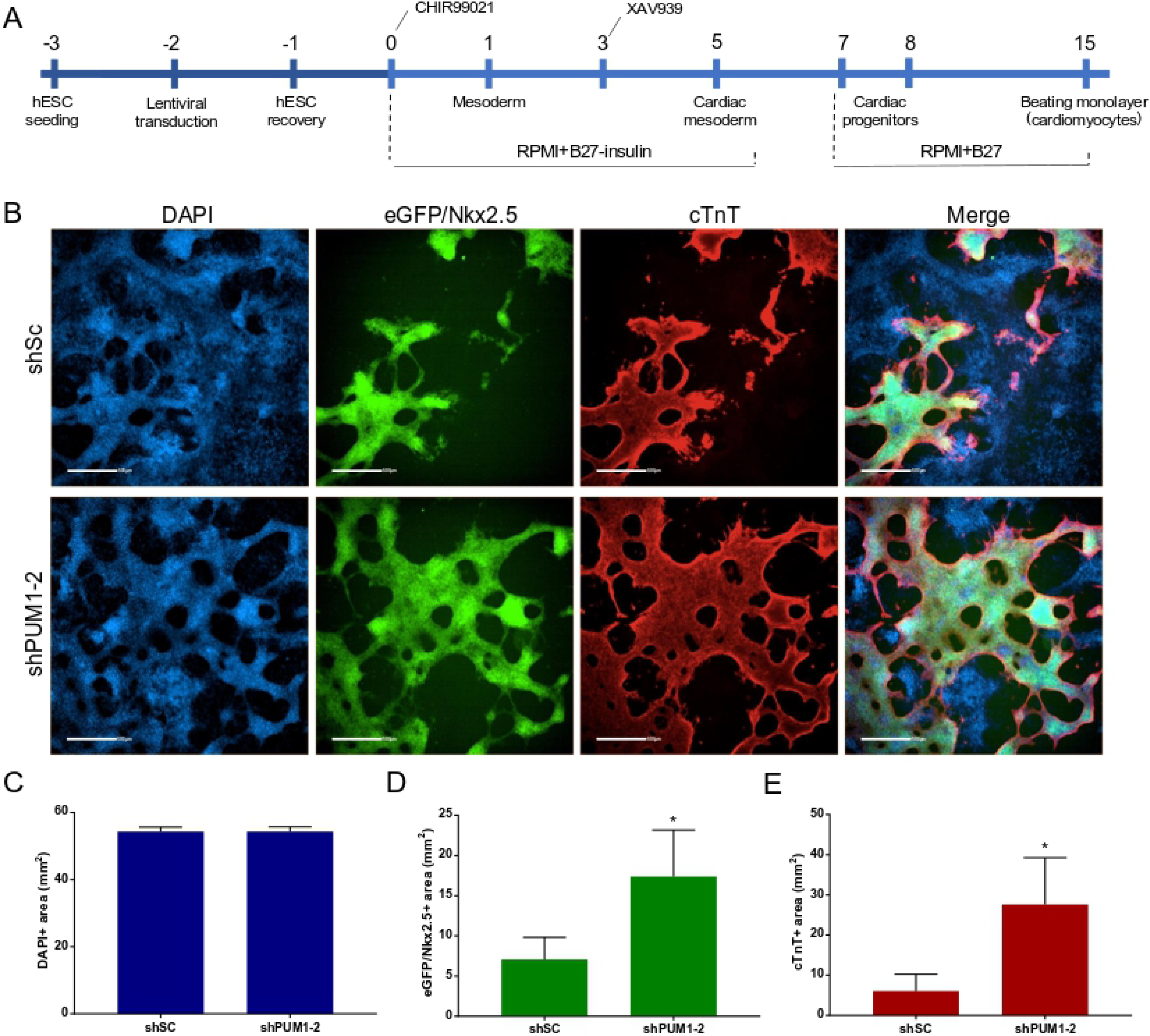
Effect of knockdown of *PUM1* and *PUM2* during monolayer cardiac differentiation. (A) Scheme of transduction and monolayer cardiomyogenic differentiation protocol of hESC. (B) Representative images of immunofluorescence of transduced hESC induced with a monolayer cardiac differentiation protocol, after 15 days. Scale bars: 500 μm. Quantification of DAPI^+^ (C), eGFP/Nkx2.5^+^ (D) and cTnT^+^ (E) staining areas after 15 days of monolayer cardiac differentiation. *p<0.05

### mRNA targets of PUM1 and PUM2 involved in cardiac development are associated with polysomes during cardiomyogenesis

Our results suggest that PUM1 and PUM2 may play a role in the cardiac differentiation process. Thus, we investigated the expression profile of the targets mRNA of PUM1 and PUM2 during cardiomyogenesis of hESCs using previously published data (32). Briefly, mRNA population associated with polysomes in five time-points (days 0, 1, 4, 9 and 15) during the *in vitro* cardiac differentiation of hESCs were isolated and sequenced (32). Using these data, we selected mRNA targets of PUM1 and PUM2 and we evaluated the genes that increase after 1 and 4 days of cardiomyogeneses (Supplementary Table S2).

Gene ontology (GO) analysis for each set of mRNAs targets of PUM1 and PUM2 upregulated in polysome were conducted with g:Profiler (37) (http://biit.cs.ut.ee/gprofiler/). The complete lists of GO analysis concerning cellular components, biological processes, molecular functions and signaling pathways could be found in the Supplementary Tables S3. In order to aid visualization of the GO terms found for biological processes we use the REVIGO (38) (http://revigo.irb.hr/). The mRNAs targets of PUM1 and PUM2 were related with biological process as stem cell development, mesenchymal cell differentiation, head development and others (Figures 5A). Networks analysis of these same genes presents elements involved with circulatory system development, among these genes are: EPOR, WNT5A, EFNA1, RHOB, HOXB3, BMP2, SMAD7, FN1, ADAMTS6, COL15A1 and COL4A2. These findings may underlie the fact that PUM1 and PUM2 silencing has favored cardiomyogenesis, since these mRNAs associated with circulatory system development may be more activated.

**Figure 5.**
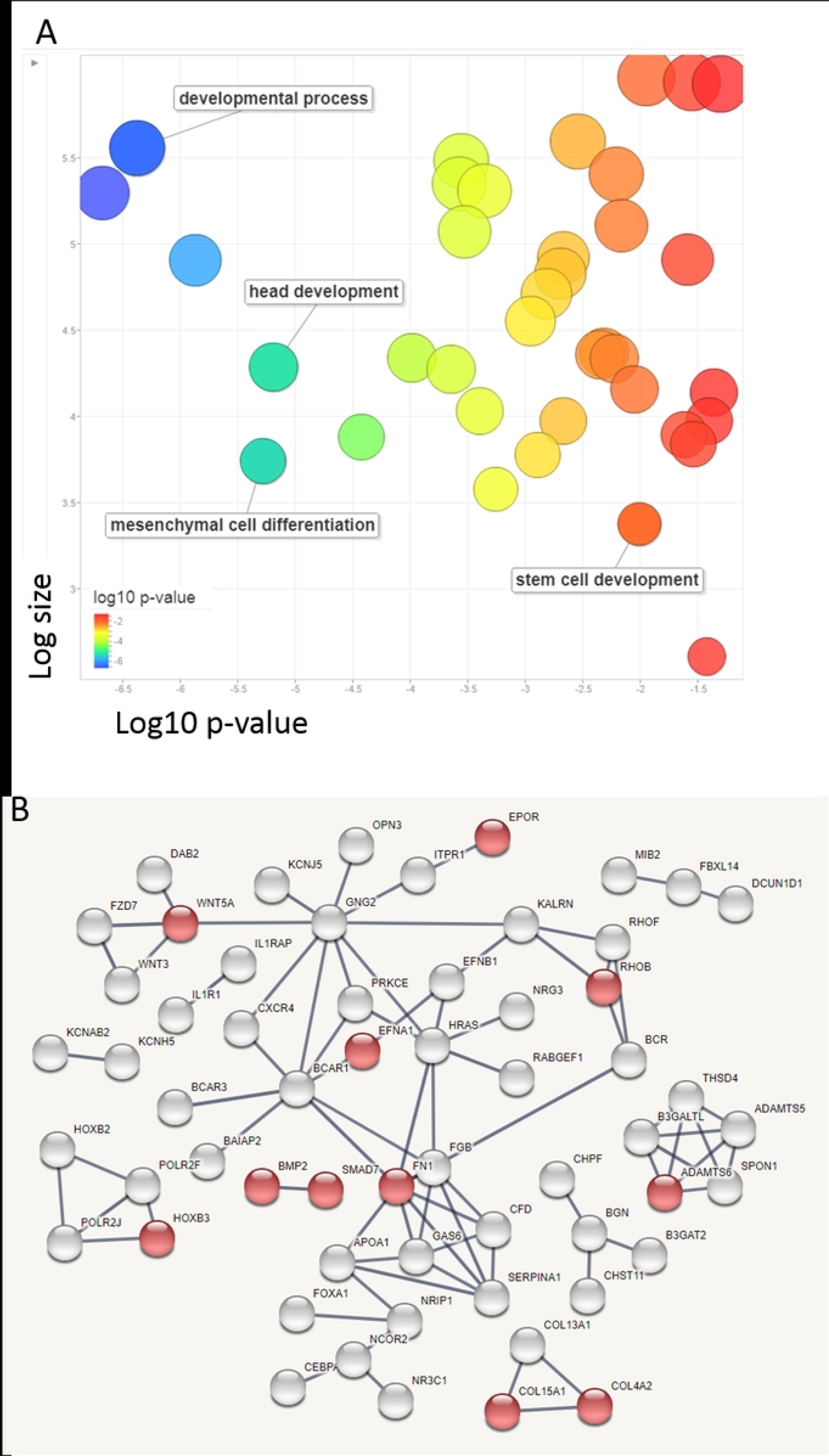
mRNA targets of PUM1 and PUM2 upregulated in the polysome during cardiomyogenesis. (A) Gene ontology analysis of mRNAs targets of PUM1 and PUM2 upregulated in 4 days of cardiomyogenesis. The figure shows a REVIGO scatterplot of the representative clusters of GO terms obtained with g:Profiler. In the two-dimensional space of the graph, the log10 p-value of each GO after REVIGO analyses is plotted on the x-axis, while the terms are scattered based on log size on the y-axis. Bubble color indicates the provided p-value (legend). (B) Networks analysis of mRNAs targets of PUM1 and PUM2 upregulated in 4 days of cardiomyogenesis. Functional enrichments in circulatory system development (FDR 1.99e-05) were represented in red. Interaction Score: Highest confidence (0.900), STRING.

## DISCUSSION

The PUF family of RNA-binding proteins is involved in these events in higher eukaryotes (40). In humans, PUM1 and PUM2 are coexpressed in several tissues, and almost 90% of the PUM2 target genes are also targets of PUM1, showing that human PUM proteins have many similarities in substrate specificity and eventually act redundantly on common targets (22,41). In this context, hESCs may help to clarify the role of these proteins, either in pluripotency maintenance or during a specific lineage differentiation.

In this work, when monitoring *PUM1* and *PUM2* expression during the cardiomyogenic differentiation of hESC, we observed that *PUM1* expression increased over time. Distinct Pumilio expression profiles were observed during mouse hematopoietic (12) and human adipogenic differentiation (42). However, when we analyzed *PUM1* and *PUM2* expression profile during cardiomyogenesis (32) we observed that both mRNAs appear associated to polysomes in a constant pattern throughout the whole process, indicating that the maintenance of *PUM1* and *PUM2* mRNA levels is important for cell function.

Therefore, we investigated the impact of Pumilio proteins on stemness and cardiomyogenesis by knocking down the two human paralogs, individually and in combination. As discussed previously, *PUM1* and *PUM2* share most of their mRNA targets (22,41), and a compensatory regulation mechanism has been observed when one of these genes is silenced by increasing the expression of the other (43). We evaluated the levels of *PUM1* and *PUM2* mRNAs after silencing these genes individually, and we did not observe this compensation, at least at the mRNA level. We hypothesized that due to their high similarity, the shRNA used to knockdown one transcript impacted the stability of the other transcript, at least in this cell type.

We observed that the *PUM1* and *PUM2* silencing altered the mRNA levels of two genes associated with the pluripotency phenotype, *OCT4* and *NANOG*. In both cases, the expression was significantly reduced in silenced cells in relation to the control, suggesting that PUM proteins may be involved in the maintenance of hESC pluripotency. These results corroborate with other studies that strongly suggest that PUF proteins may mediate a widespread and ancient mechanism for repressing the differentiation and maintaining the self-renewal of stem cells (10,15,44–47).

As Pumilio proteins seem to regulate the stemness phenotype, we examined the effect of their silencing on hESC induced to two different protocols of *in vitro* cardiomyogenesis. In EBs protocol, hESC silenced for *PUM1* and/or *PUM2* showed an increase in the percentage of cTnT^+^ cells, despite it was not statistically significant. However, it is important to point out that the percentage of positive cells during EBs cardiac differentiation protocol may even be higher, since the dissociation of EBs containing organized cell-cell junctions becomes difficult, which may impair quantification by flow cytometry. Moreover, using the monolayer protocol we observed that *PUM1-2* silenced hESC exhibited a larger cTnT staining area than control cells. These changes indicated that the reduction of *PUM1* and *PUM2*, even over a short period, can increase the number of cardiomyocytes generated from *in vitro* cardiac differentiation protocols. The mechanism behind these effect needs to be investigated but could be related to the decrease of pluripotent markers.

Because stemness and differentiation are both subjected to Pumilio regulation, based on the data previously generated from mRNA sequencing from free and polysomal fractions of hESC induced to cardiomyogenesis (32), we performed a bioinformatic analysis of the expression profiles of the PUM targets. We identified two clusters of genes consistently increasing their representation during differentiation. GO analysis revealed that genes related to nervous system development were present in the free fraction, while in the polysomal fractions we identified more genes related to mesodermal and cardiac development. This evidence suggested that mRNAs associated with other germ layers were enriched in the free fraction and, possibly, are being switched off during the cardiac differentiation process. On the other hand, genes related to cardiomyogenic lineage are associated with polysomes, suggesting, as expected, that these genes are expressed and translated. Besides that, among Pumilios targets mRNAs we identified many transcription factors, which are key components of differentiation processes. These results suggest that PUM1 and PUM2 could be regulators of transcription factors expression during cardiomyogenesis.

Here we demonstrated that *PUM1* and *PUM2* are expressed throughout of hESC *in vitro* cardiomyogenesis differentiation. Moreover, when both genes were knocked down, we observed a reduction in pluripotency transcription factors expression and an increase in cardiac differentiation efficiency. Through analysis of previous published RNAseq data, we could identify many PUM targets mRNA modulated during cardiomyogenesis. Despite that, the mechanism of these proteins in cardiac lineage commitment still needs to be elucidated.

## ADDITIONAL INFORMATION

### Ethics approval and consent to participate

Not applicable.

### Competing interests

The authors report there are no potential conflicts of interest or financial interests.

### Funding

This study was supported by FIOCRUZ.

### Authors’ contributions

ILZS and AWR carried out the molecular studies. GCC and LS carried out the bioinformatics analysis. AWR and MS carried out the cytometry analysis. BD review the manuscript. DFG and PS wrote the manuscript. All authors read and approved the final manuscript.

## Acknowledgments

The authors thank the Program for Technological Development in Tools for Health-PDTIS/FIOCRUZ for the use of its facilities. We also thank Wagner Nagib de Souza Birbeire for editing the videos.

